## Supplementary Figures for "Automatic mapping of multiplexed social receptive fields by deep learning and GPU-accelerated 3D videography"

### Supplementary Figures and Legends

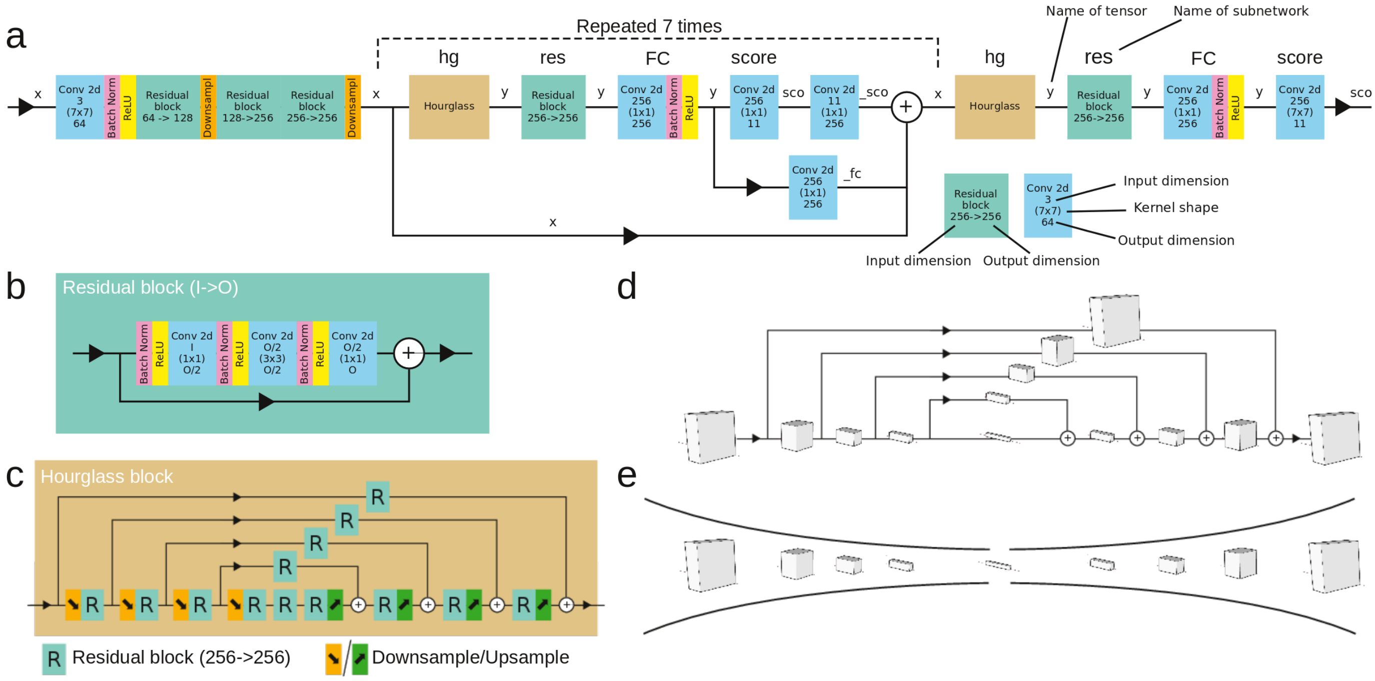


**Supplementary Figure 1. Deep convolutional neural network architecture. a**, Schematic of flow of tensors through deep convolutional neural network. Convolutional blocks show kernel shapes and input/output dimensions of feature dimension, starting from 3 (RGB image), expanding to 256 during hourglass blocks, and ending in 11 for intermediate and final outputs (4 body point targets, 7-part affinity fields). In the drawing, the hourglass stack is repeated 7 times (the default number), but we find that repeating only 3 times is sufficient for our purpose (see following supplementary figures). Full implementation details (e.g., including stride, padding, bias, etc.) are included in supplementary code. **b**, Schematic of a single residual block. **c**, Schematic of a single hourglass block. Upsampling (green, nearest neighbor) and downsampling (orange, by max pooling, both a factor of 2) happens along 2D image space (height/width). **d**, Shapes of tensors flowing through hourglass block. Along bottom path, feature dimension stays constant, but image dimensions (height/width) are increasingly downsampled, and then upsampled again. After each upsample, tensors are merged with skip connections (paths above). **e**, The hourglass-like shape that gives name to the network architecture.

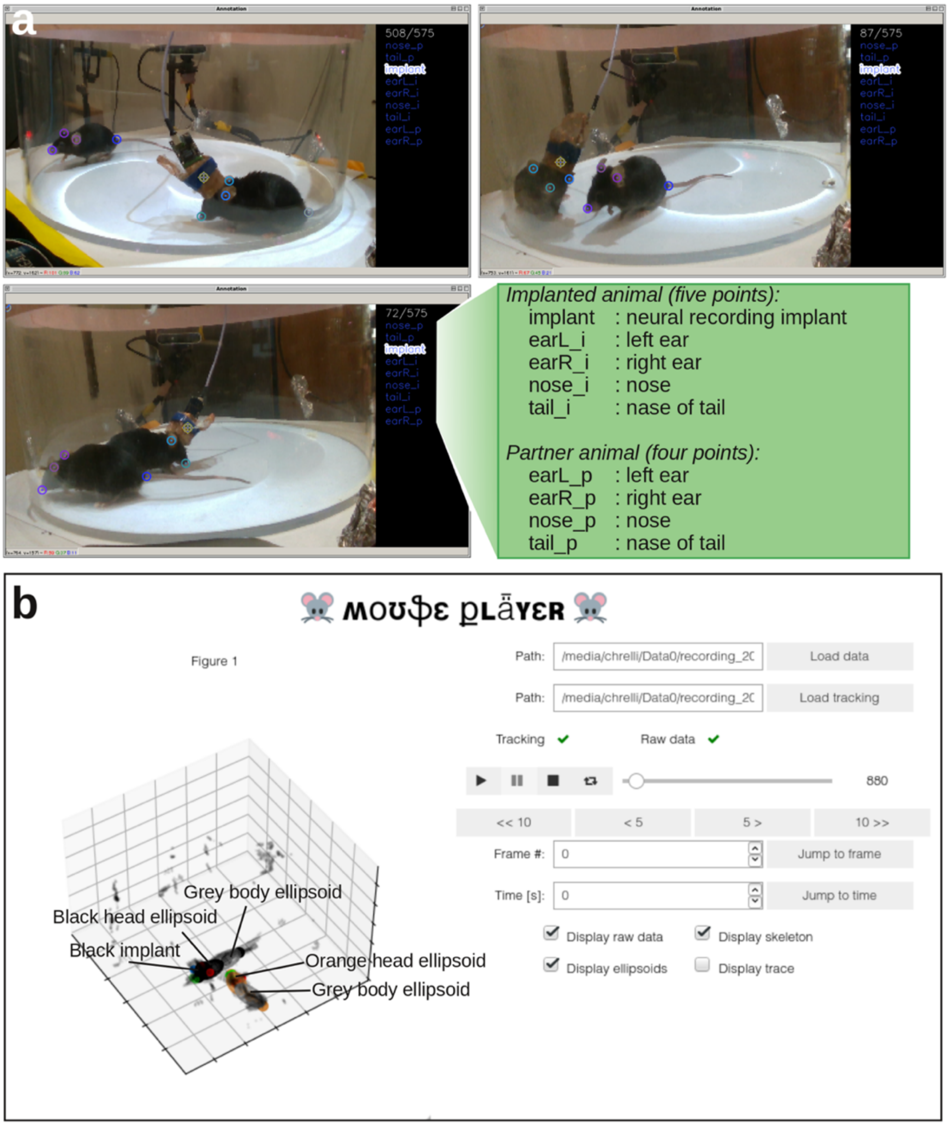


**Supplementary Figure 2. GUIs for labeling of training data for the neural network and for viewing tracked data. a**, For training the network to recognize body parts, we must generate labeled frames by manual annotation. For each frame, 1-5 body parts are labeled on the implanted animal and 1-4 body parts on the partner animal. This can be done with any annotation software; we used a modified version of the free 'DeepPoseKit-Annotator' (Graving et al., 2019) (https://github.com/jgraving/DeepPoseKit-Annotator/) included in the supplementary code. This software allows easy labeling of the necessary points, and pre-packages training data for use in our training pipeline. Body parts are indexed by i/p for implanted/partner animal ('nose_p' is the nose of the partner animal, for example). **b,** GUI for viewing and quality control of tracked behavior (raw data, body skeleton, ellipsoid surfaces and time trajectory) running in an interactive Jupyter notebook.

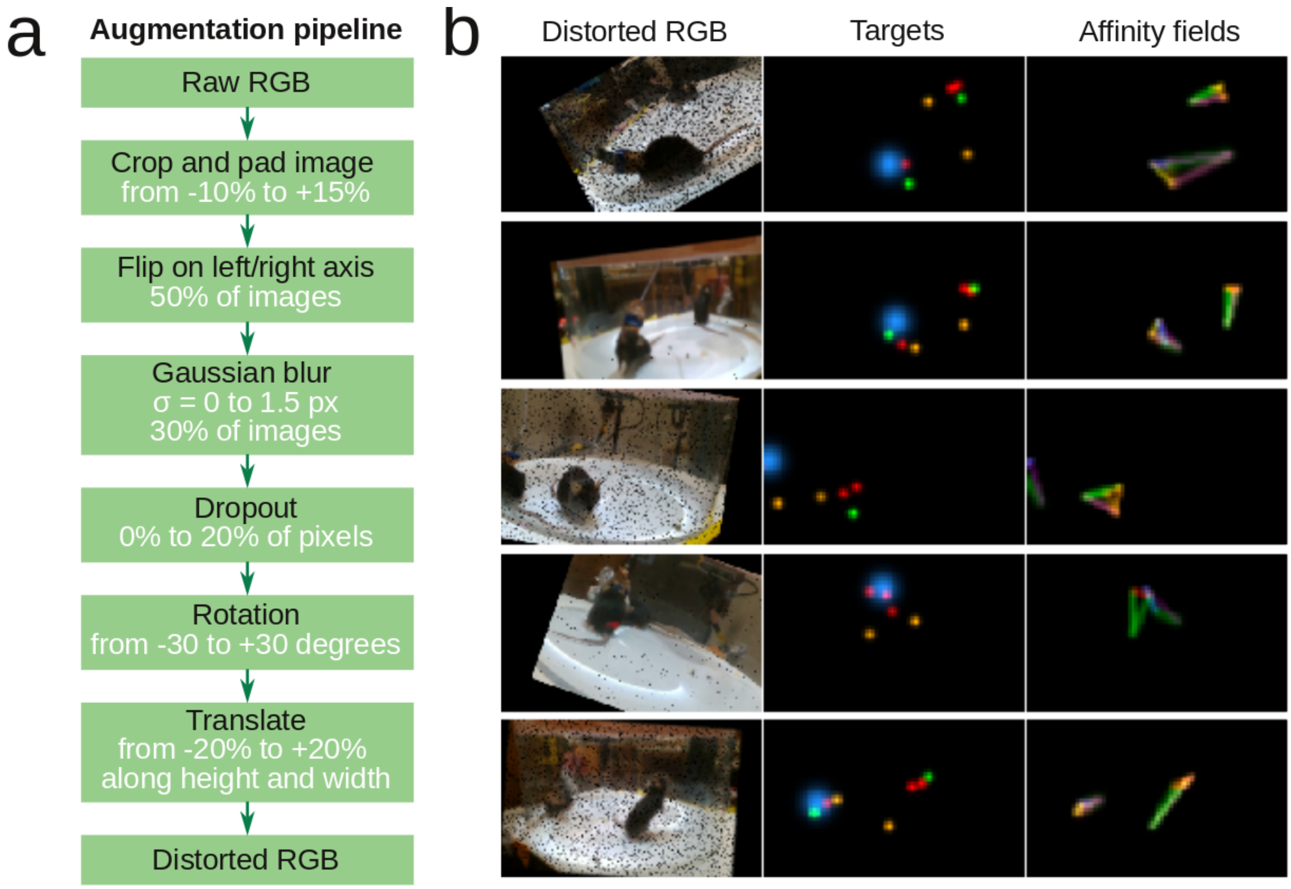


**Supplementary Figure 3. Augmentation pipeline for network training. a**, Flowchart of augmentation pipeline used to generate distorted frames during network training. **b**, Examples of distorted labeling frames generated by augmentation pipeline, as well as corresponding body part targets and affinity fields used during training.

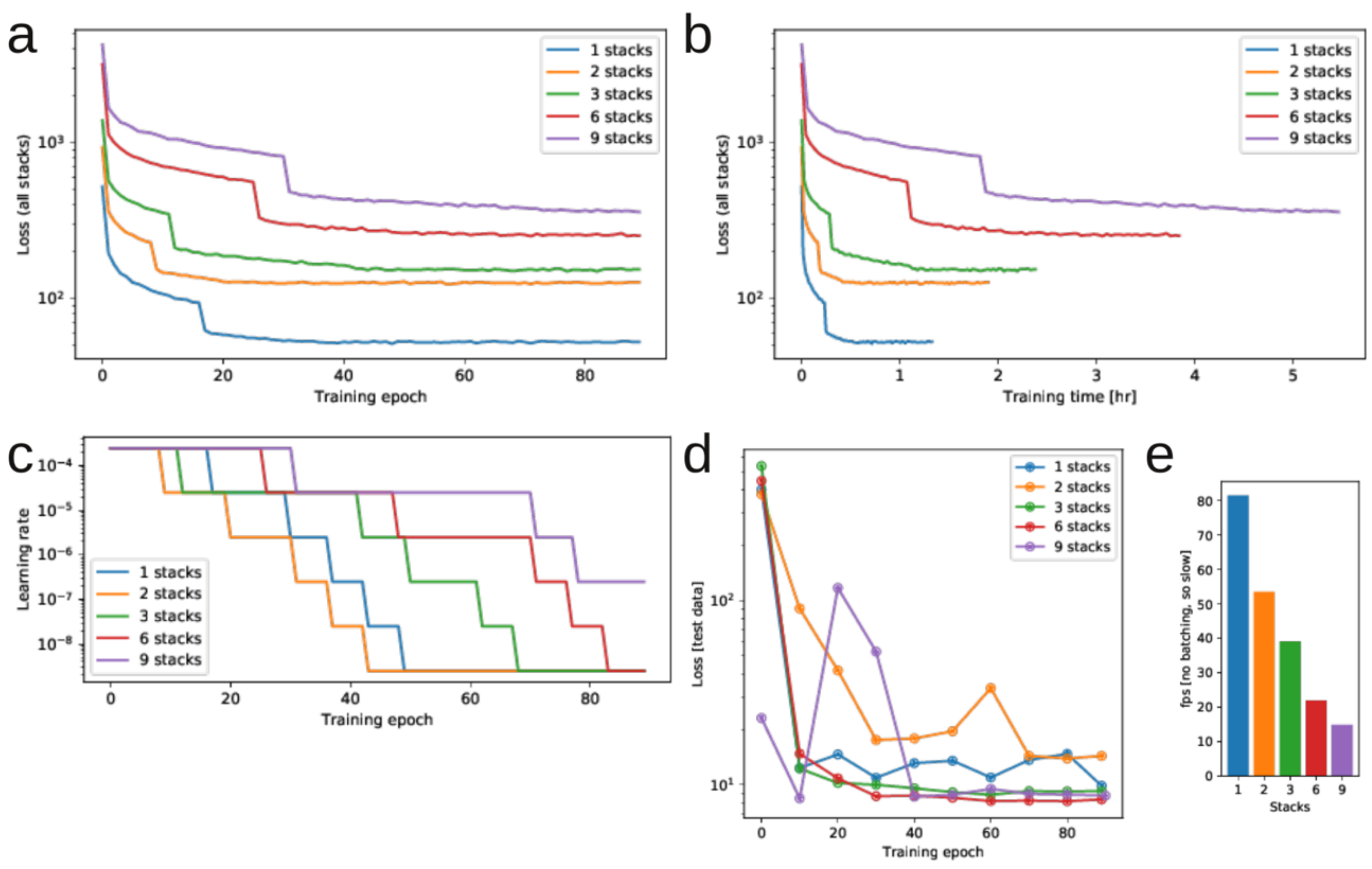


**Supplementary Figure 4. Network training history and performance as a function of hourglass stacks. a**, Loss history as a function of training epoch, for networks with 1, 2, 3, 6, and 9 hourglass stacks. **b,** same as panel a, but as a function of training time. **c,** Learning rate schedule (automatically adjusted) as a function of training epoch. **d,** Test loss, as a function of training epoch, for networks with 1, 2, 3, 6, and 9 hourglass stacks. Beyond 3 stacks, there was little improvement in the training loss. **e,** Relative inference time with no batching (since the batch size will have to be smaller for a network with more stacks – for real use, we used batched inference).

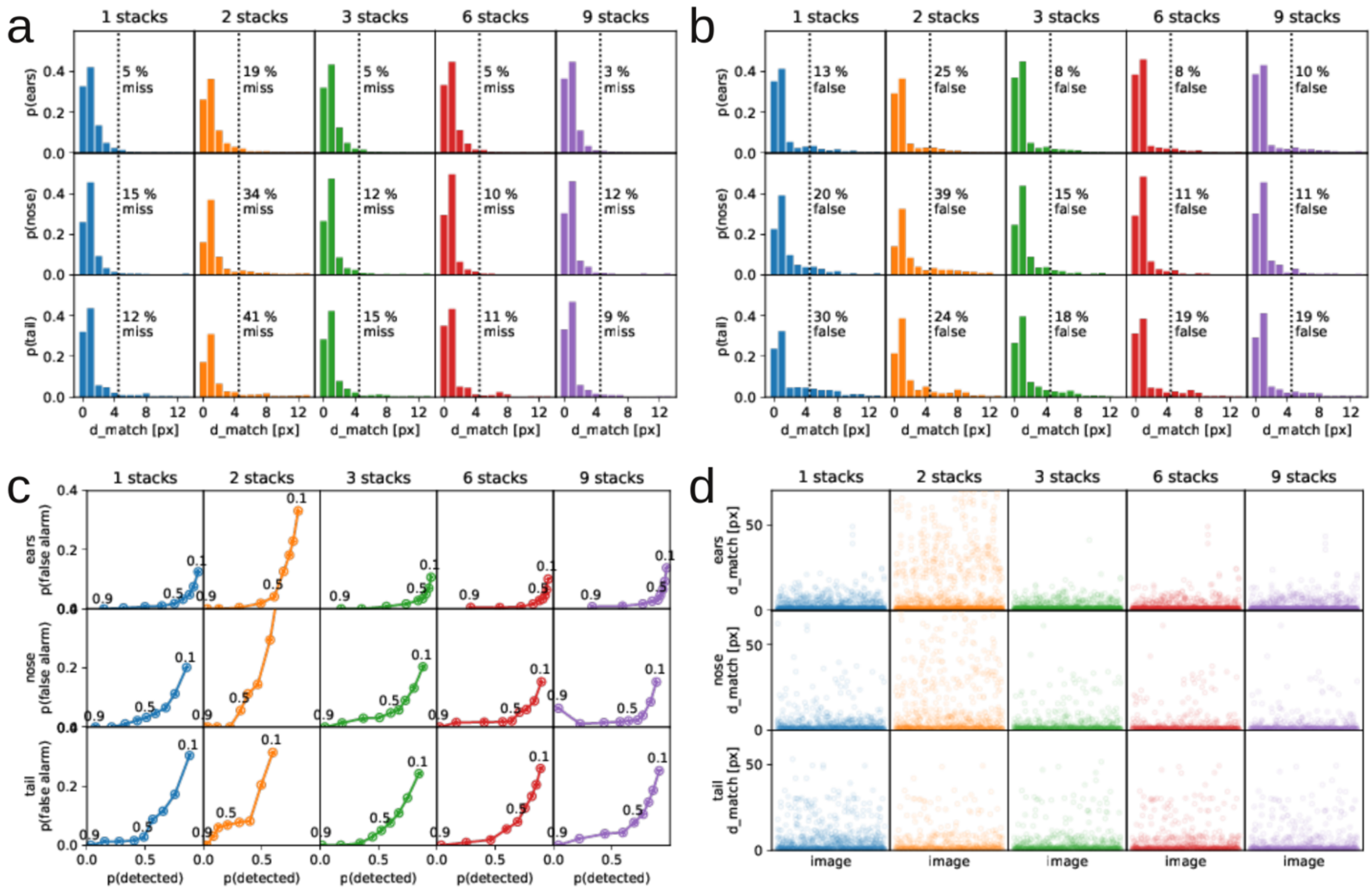


**Supplementary Figure 5. Network sensitivity and precision as a function of hourglass stacks. a,** Number of 'missed' keypoints, as a function of number of network stacks, after full training, for ear, nose and tail keypoints (the three rows). We count a hand-labeled body keypoint as found by the network as 'detected' if it was within 5 pixels of a keypoint suggested by the network. **b,** Same as **a**, but counting the number of 'false detections'. We defined a false detection, as a keypoint suggested by the network, which was more than 5 pixels from a corresponding hand-labeled keypoint. **c,** Plots of false alarm rate (p(false alarm)) as a function of detected rate (p(detected)) for all keypoints in the test data, plotted for different probability cutoffs (from 0.1 to 0.9 in 0.1 steps, indicated by dots on the curves). **d,** Shortest distance from a proposed keypoint location to a hand-labeled keypoint location (d_match), across images in the test data, for ear, nose and tail keypoints (rows) as a function of image stacks (columns).

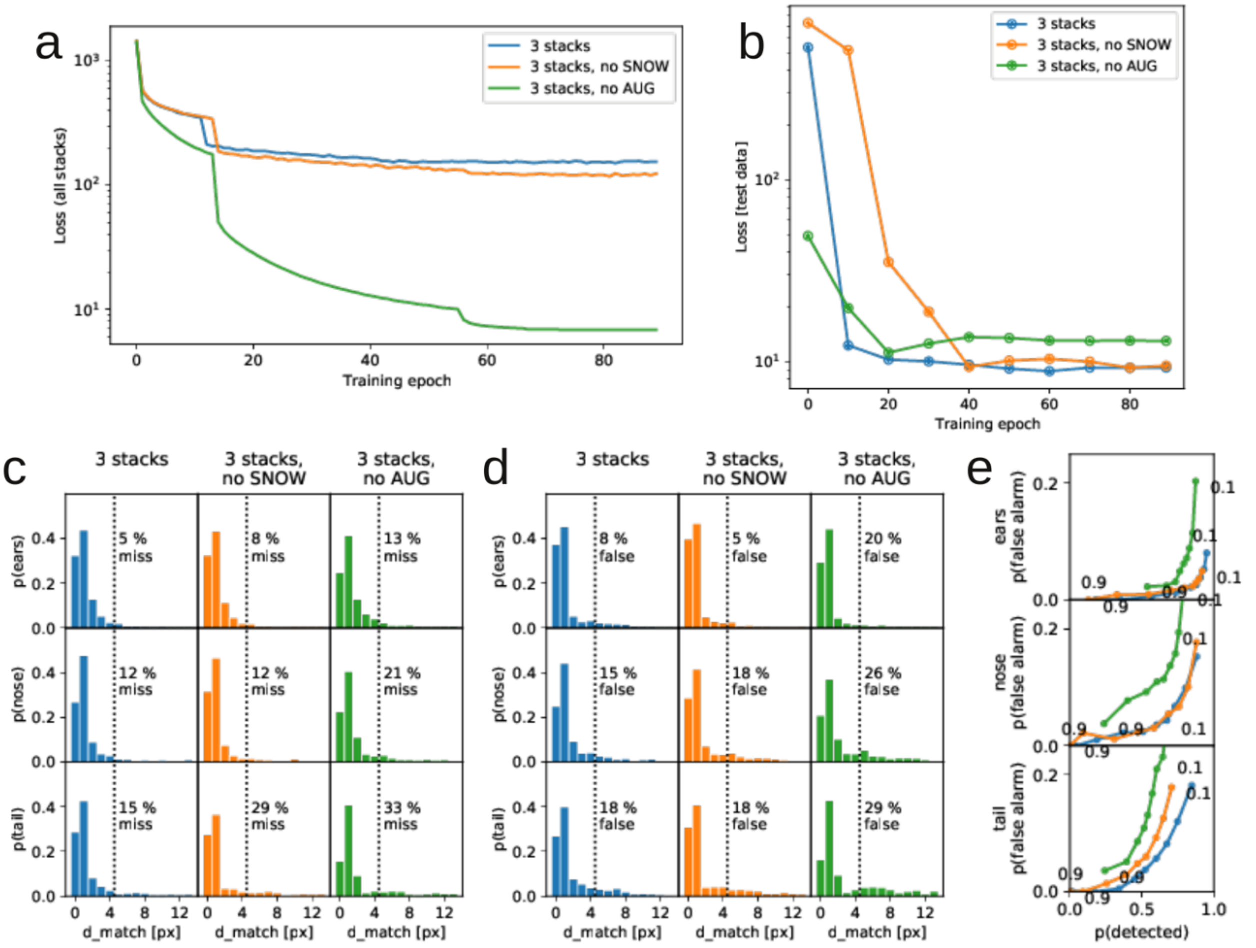


**Supplementary Figure 6. Performance of the network, with and without different types of augmentation. a**, Training loss as a function of training epoch, for networks with all augmentation ('3 stacks'), without the artificially generated laser dot pattern ('no SNOW'), and without any augmentation ('no AUG'). **b,** Loss on the test data, as a function of training epoch. **c,** Percentage of missed keypoints, with/without different levels of augmentation. **d,** Percentage of falsely detected keypoints, with/without different levels of augmentation. **e,** Plots of false alarm rate (p(false alarm)) as a function of detected rate (p(detected)) for all keypoints in the test data, plotted for different probability cutoffs (from 0.1 to 0.9 in 0.1 steps, indicated by dots on the curves), across the three levels of augmentation. The artificially-generated laser dot pattern does not generally improve detection of ears and noses, but makes a substantial difference in the detection of the tail keypoints.

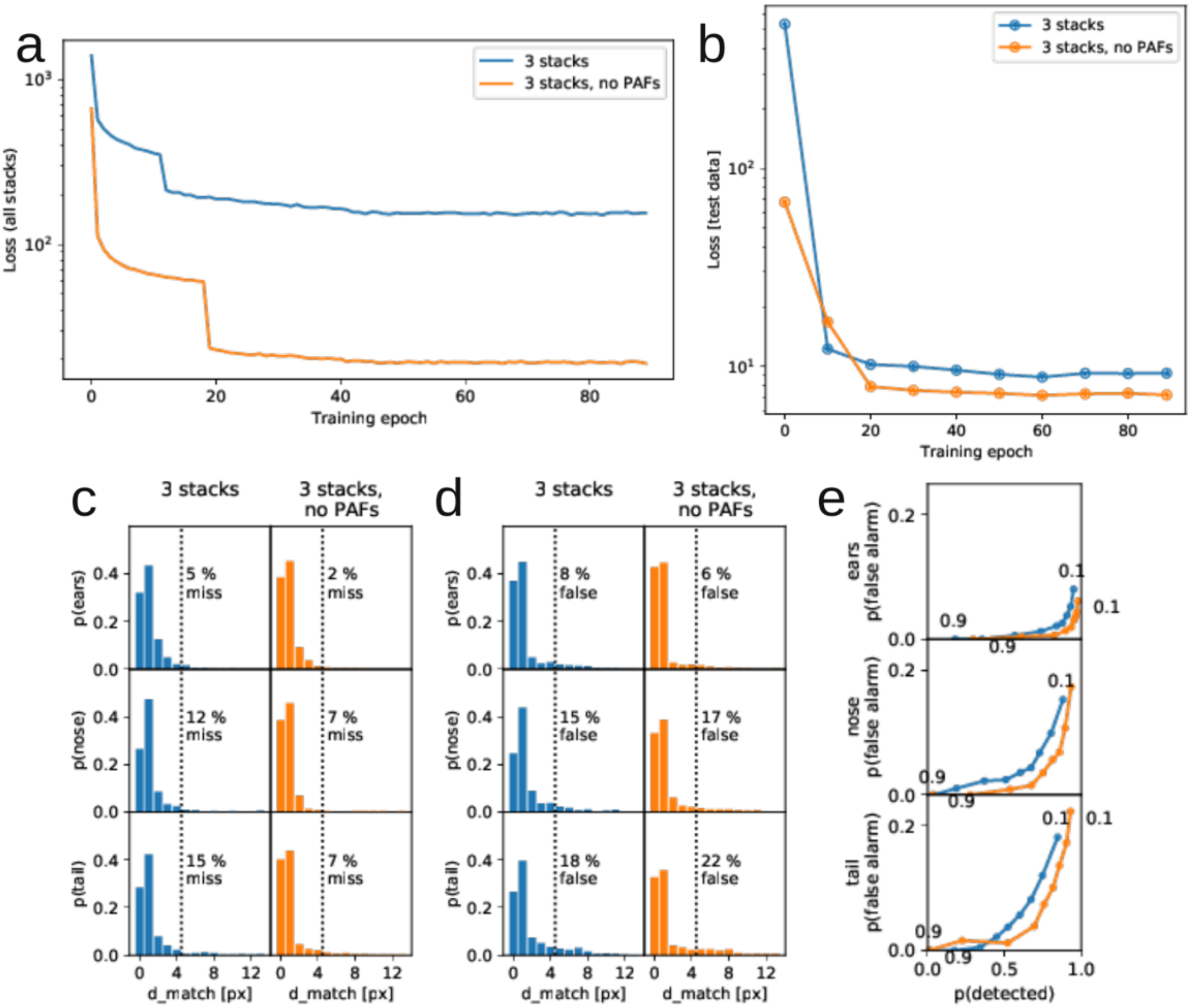


**Supplementary Figure 7. Performance of the network, with and without part affinity fields (PAFs).** **a**, Training loss as a function of training epoch, for networks with/without PAFs. **b,** Loss on the test data, as a function of training epoch. **c,** Percentage of missed keypoints, with/without PAFs. **d,** Percentage of falsely detected keypoints, with/without PAFs. **e,** Plots of false alarm rate (p(false alarm)) as a function of detected rate (p(detected)) for all keypoints in the test data, plotted for different probability cutoffs (from 0.1 to 0.9 in 0.1 steps, indicated by dots on the curves), with/without PAFs. Note that contrary to expectation, inclusion of PAFs usually did not improve the keypoint detection performance, over a network with an identical dimensionality but where the network was not required to learn PAF representation. The network without PAFs had a similar architecture to the network with PAFs. The representation without PAFs led to a lower overall loss and a lower number of false alarm and missed detections. As the performance differences with/without forcing the PAFs are minimal, our code imposes the PAFs by default.

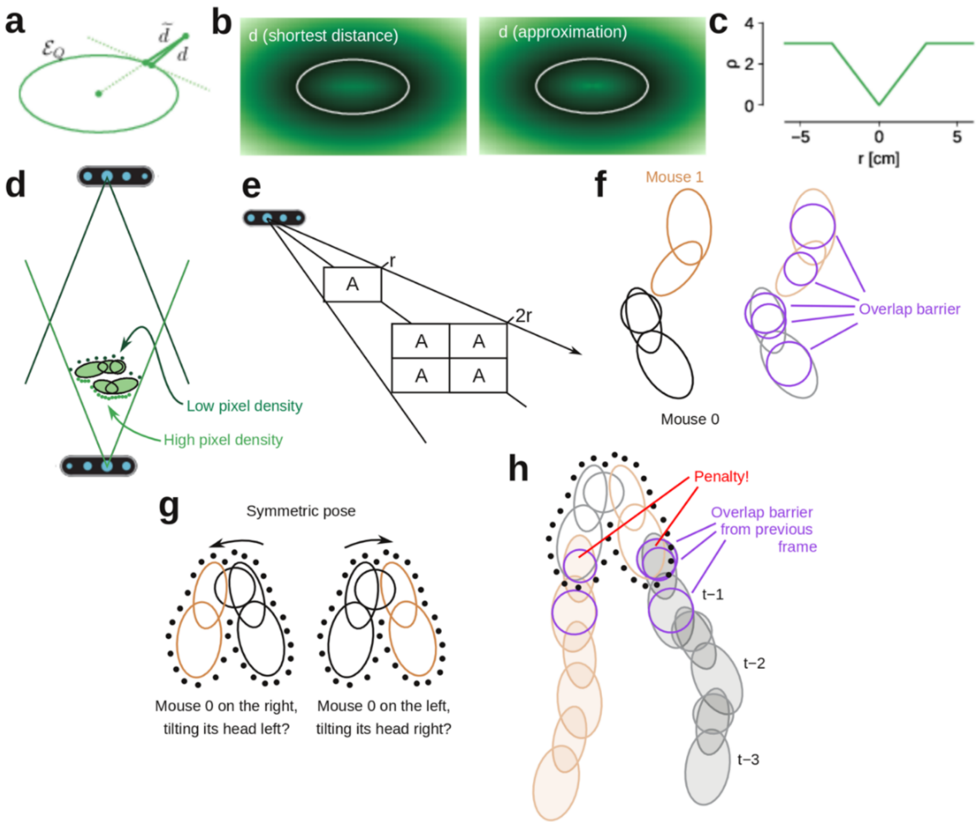


**Supplementary Figure 8. Loss function calculation details. a**, Shortest distance to surface of an ellipsoid, $d$, and our approximation, $\tilde{d}$. **b**, $\tilde{d}$ is a good approximation to $d$. Color map, value of $d/\tilde{d}$. White line, ellipsoid surface. **c**, The loss function, ρ, associated with the pointcloud is the mean absolute error of the distance estimate, truncated at +/- 3.0 cm **d**, Pixel density of the point-cloud depends on distance to the fixed-resolution depth cameras. **e**, Pixel density is inversely proportional to the square of the distance to depth camera. **f**, Overlap barrier spheres (implant sphere and spheres centered on the body ellipsoids with a radius equal to the minor axis). **g**, Example of mirror symmetric body position (side-by-side, facing same direction), resulting in ambiguity in animal identity if only one frame is considered. **h**, To include the context of previous frames, we add an overlap loss penalty (similar to **f**) between each mouse and the position of the interaction partner in the previous frame. In panel **g**, right, we would add a penalty term to the particle representing joint body pose. In contrast, in panel **g**, left, this penalty is zero as there is no overlap with the position of the conspecific in the previous frame.

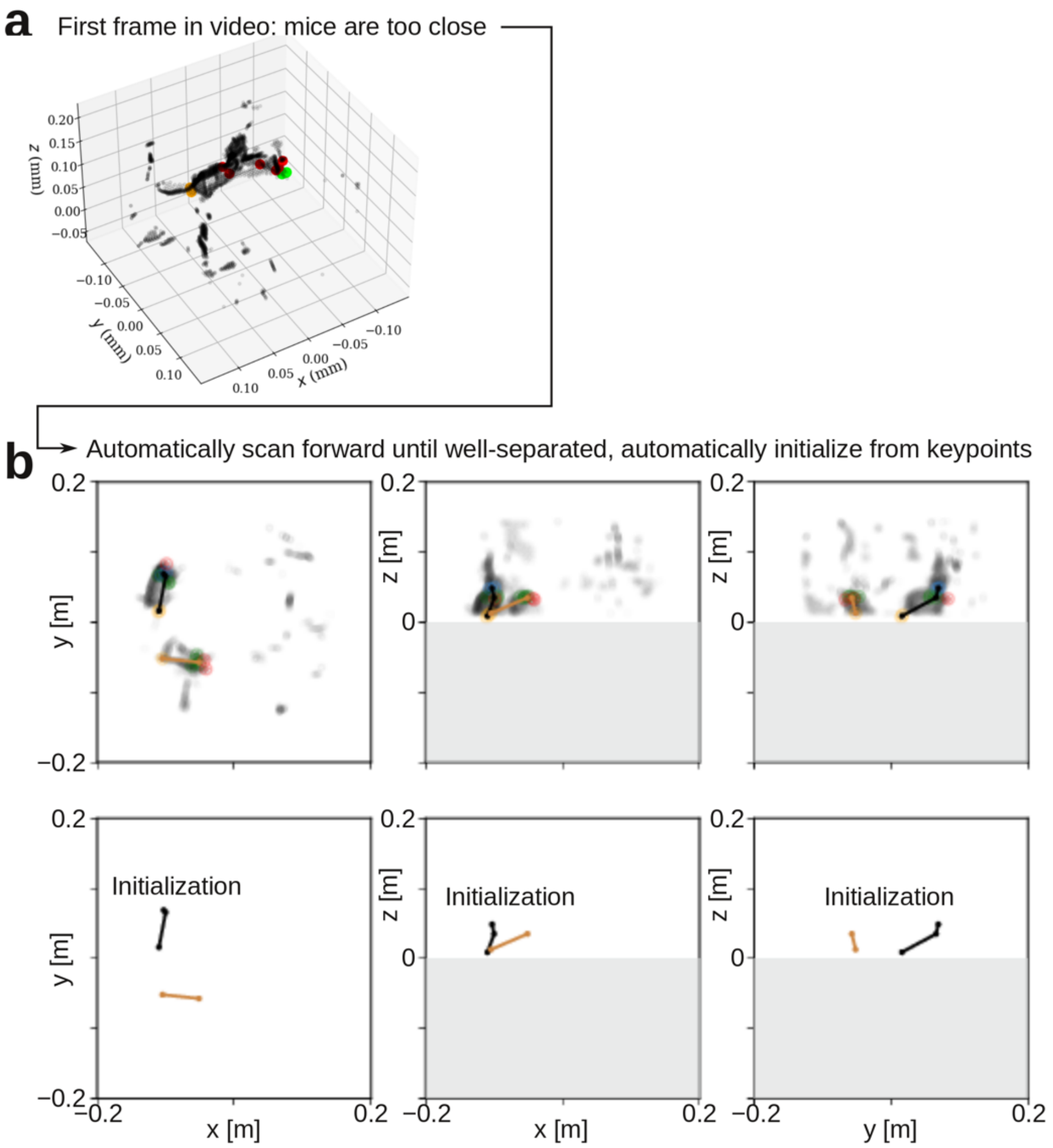


**Supplementary Figure 9. Automatic initialization procedure for the tracking algorithm. a**, Example starting frame, where the mice are too close together for the automatic initialization. **b**, By scanning forward in the 3D video, the algorithm finds a frame, where both the "head cluster" (cluster of detected ear/nose points) and the "tail cluster" (cluster of detected tail points) are separated by a threshold distance. The algorithm uses an average of the head, tail and implant clusters to initialize the tracking procedure (initial guess shown by the black/brown lines, top row of plots shows the pointcloud data, bottom row of points shows only the initial guesses).

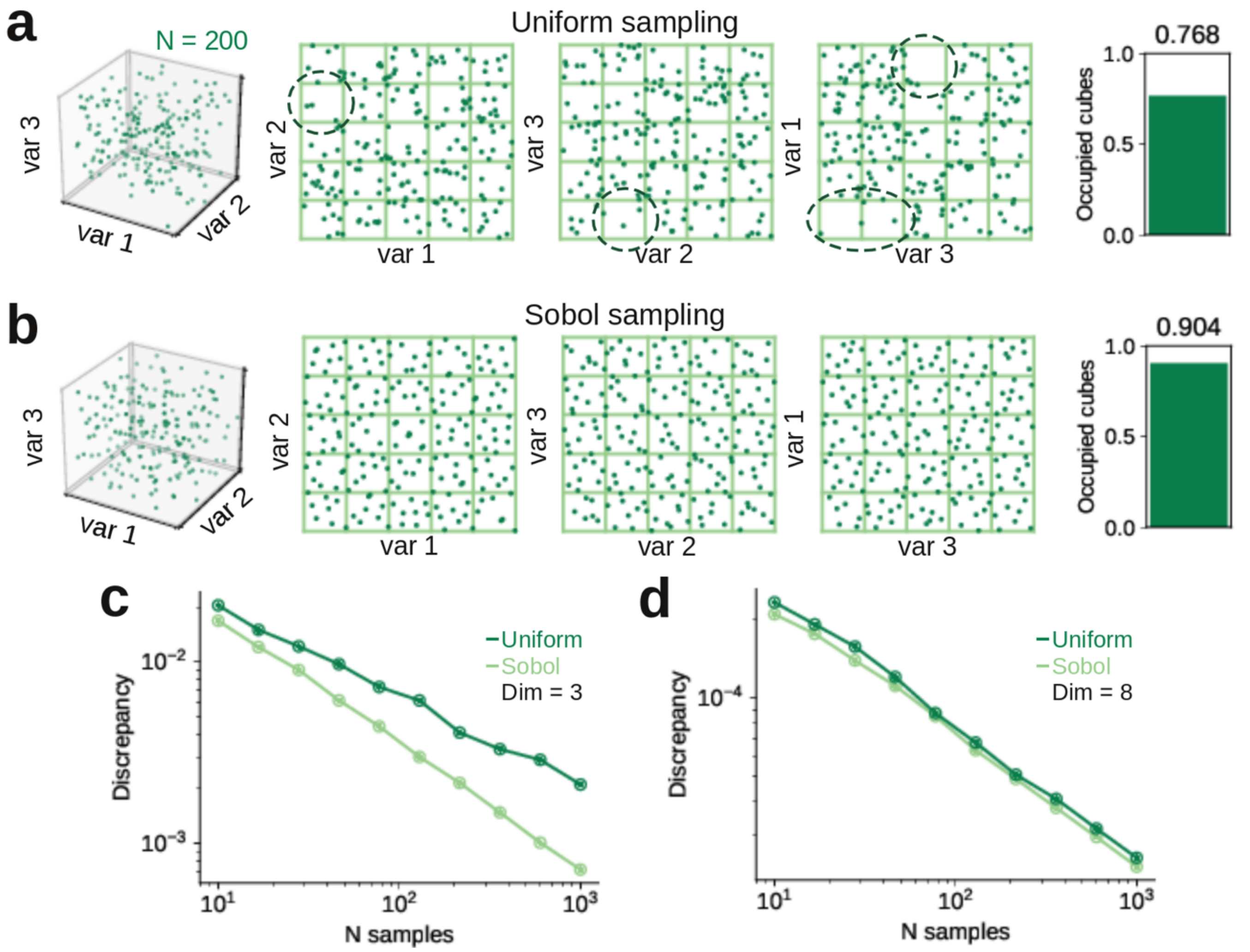


**Supplementary Figure 10. Quasi-random particle filter exploration strategy. a**, Left, 3D plot. Middle, 2D projection plots of three random variables, drawn from independent uniform distributions. Points in 3D space do not fill space well; in the 2D projections, there are squares (i.e., full rows, columns and pipes) of the 3D space not sampled at all (dashed lines). Right, partitioning space in 20%-cubes (green lines), only 76.8 % of cubes are occupied. **b**, Same as **a**, but variables are drawn from quasi-random Sobol sequence (Sobol, 1967). Points are more evenly dispersed in space, and 90.4% of all 20%-cubes are sampled. **c**, Mean discrepancy as function of sample number, for 3-dimensional (like panels **a,b**) uniform random sequence and a Sobol sequence, calculated across 100 random sequences. The Sobol sequences have a lower discrepancy, i.e. sample more regions of space. **d**, Same as **c**, but for 17-dimensional variables (like our joint body posture particles).

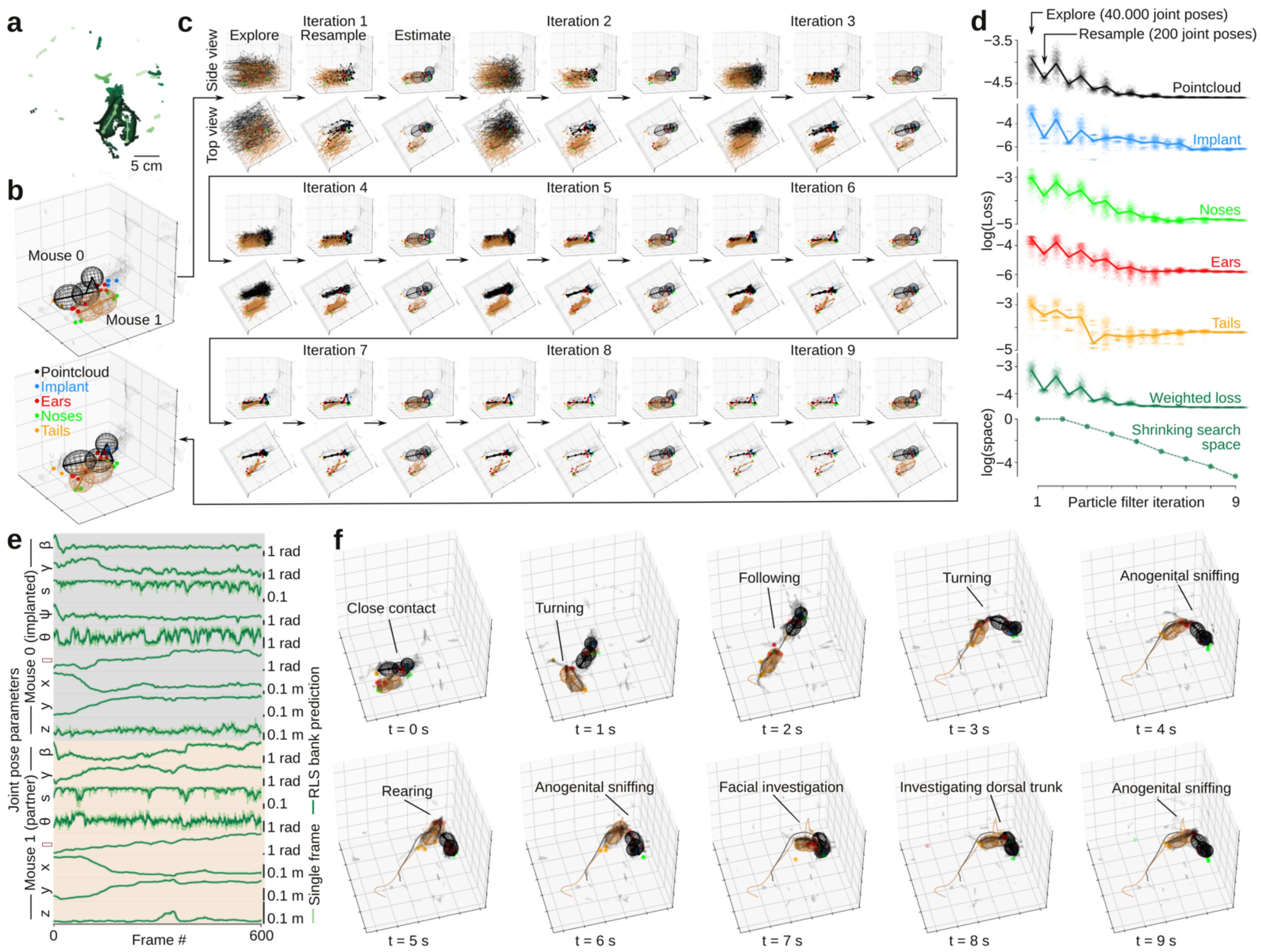


**Supplementary Figure 11. Particle filter convergence and examples of tracked behavioral data. a,** Example of manual initialization of the tracking algorithm, by manual clicking of approximate locations of the two animals (light green dots, lines) on a top-down view of the behavioral arena (dark green dots, shade indicates z-coordinate). **b,** 3D view of initialized body model (top) and fitted body model (bottom) after running tracking algorithm on the frame. Black wireframe model, implanted mouse; brown wireframe model, partner animal. **c,** Particle filter state across 9 iterations of the fitting algorithm. After iteration 2, we shrink ('anneal') the exploration space with each step. **d,** Loss function values and size of filter search space across filter iterations. **e**, Tracked data (light green) and running adaptive estimate (dark green) across 600 frames (10 s). **f**, Data and fitted joint posture model, across 10 seconds of behavior. Trailing lines, location of hip ellipsoid center in the last 10 seconds.

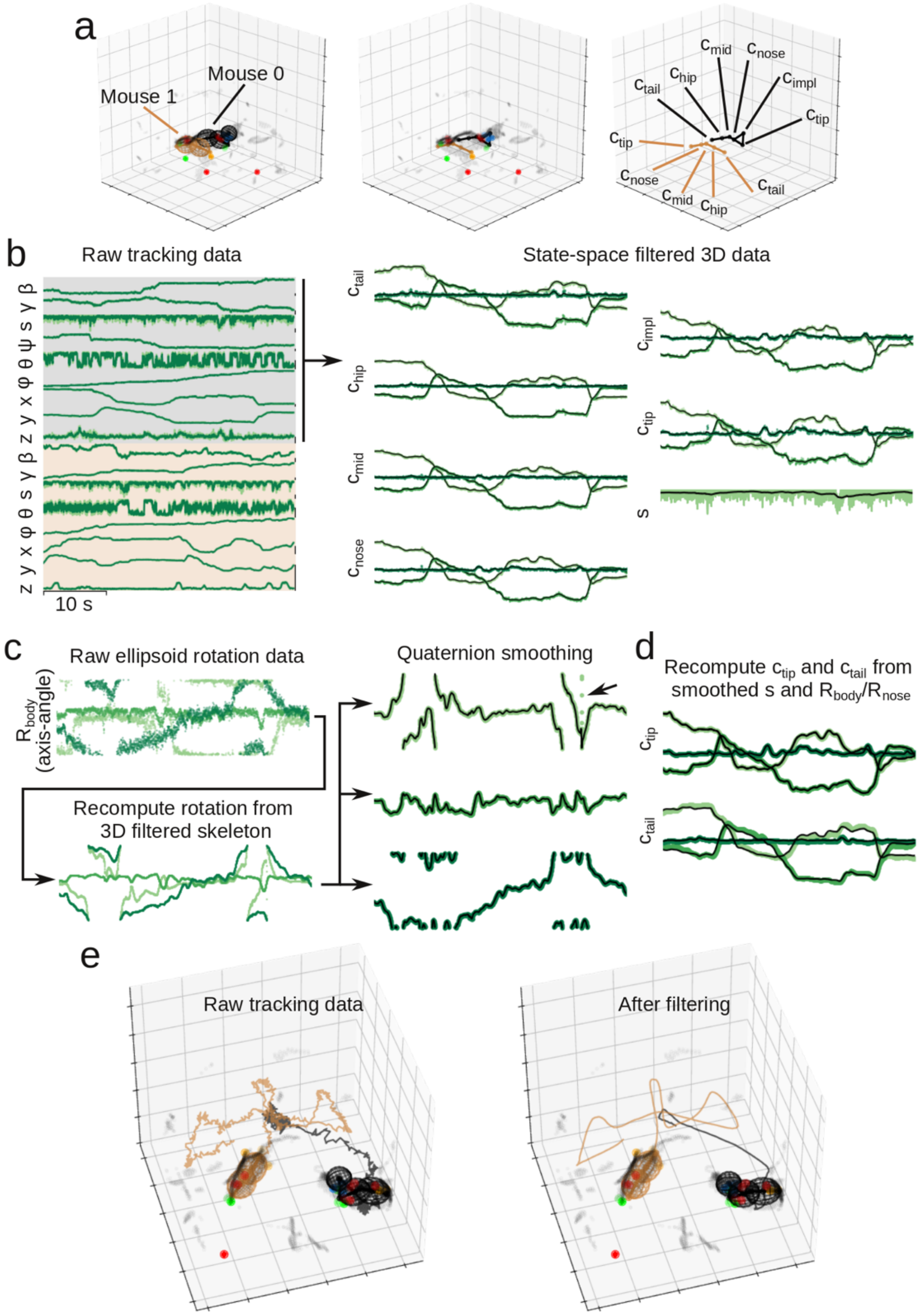


**Supplementary Figure 12. State space filtering of tracked body models. a**, Estimated 3D locations of body model surfaces (wireframes, left) and skeletons (dots and lines, right) for an example frame. **b**, Fitted joint pose parameters for the two mouse body models (left, 100 s snippet) and corresponding 3D coordinates of the body skeleton points, and the spine scaling, *s*, for the implanted mouse (right, same 100 s). **c**,

Raw 3D rotation angle of the nose ellipsoid of the implanted animal (axis-angle representation), recalculated 3D rotation angles from the filtered skeleton points, and final 3D rotation angles after quaternion smoothing (note the smoothing out of noise, indicated by arrow). **d**, Recalculated c_nose and c_tail from the smoothed 3D rotations and smoothed spine scaling. **e**, Example frame before (left) and after state space filtering of the tracked data (right).

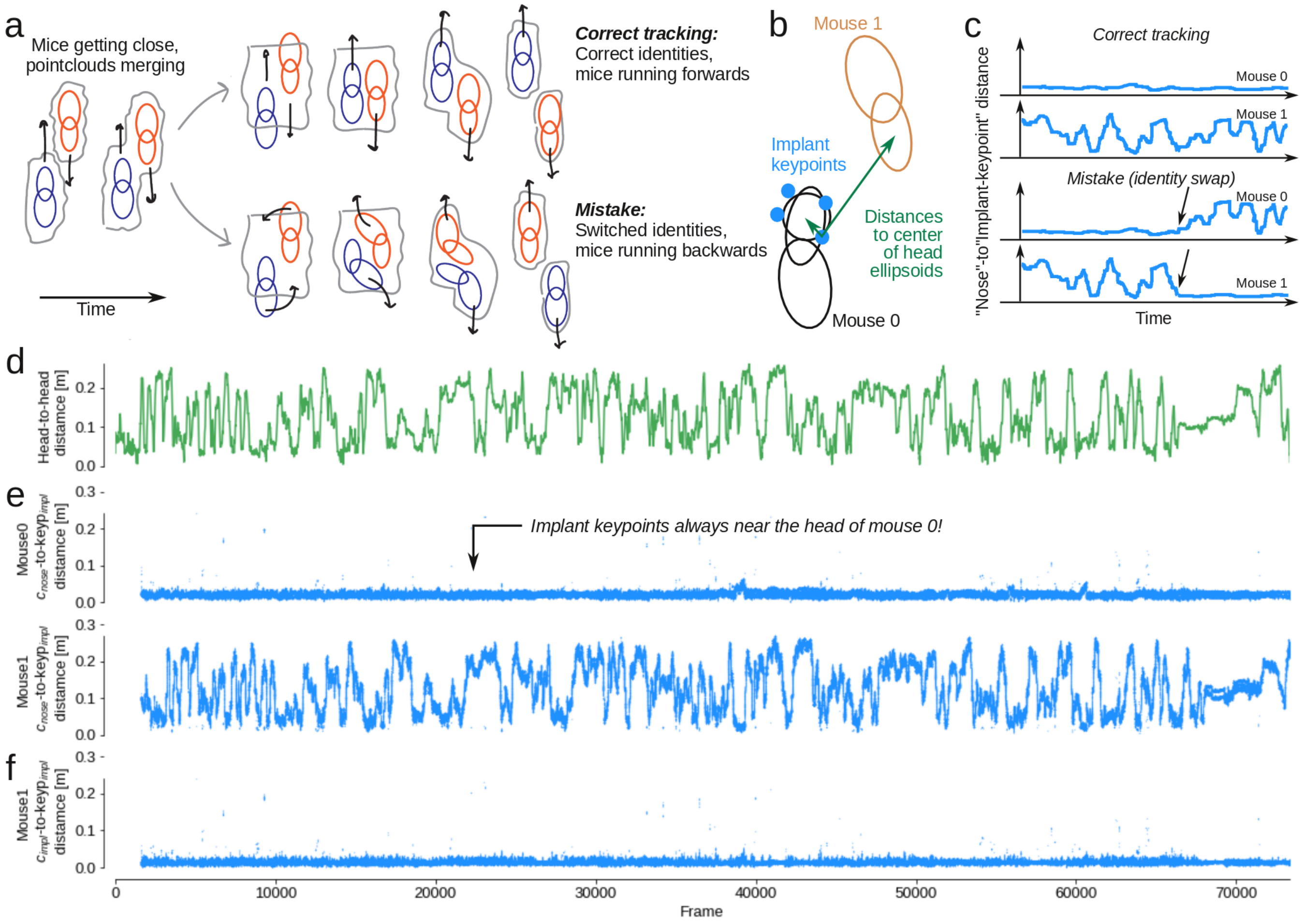


**Supplementary Figure 13. Implant-to-nose distance demonstrates that there are no swapped identities. a**, Schematic showing two common errors in tracking algorithms: Swapped identities and swapped directions. When the mice approach each other, their point clouds will merge. Because resolution and frame rates are limited, it can be hard to estimate body postures in this configuration. For example, if the tracking algorithm is not properly spatiotemporally regularized, the algorithm might mistakenly swap mouse identities, such that the mice appear to be running backwards (shown in bottom row). Direction swaps and identity swaps can also happen independently. For example, when mice are allogrooming, or passing over/under each other, identities might swap, but both mice can still appear to run normally with no apparent errors. Conversely, when a mouse is self-grooming, their point-cloud essentially resembles a ball, and when they start moving again, it may not be clear if they are 'really' moving forward or backwards. **b**, For all frames, we calculated the distance between implant key-points and the centroid of both nose ellipsoids. **c**, If there is an identity swap of the mice, this is will be evident in the distance between the implant key-points and the head of both mice. In correct tracking (top row), implant body model always follows the same mouse. In tracking with mistakes (bottom row), implant will switch from being close to one mouse, to being close to the other mouse. **d**, The head-to-head (nose-centroid-to-nose-centroid) distance for the two mice, across the session. The mice often closely interact (low head-to-head distance), allowing for potential identity swaps. **e**, Distance between implant key-points and the nose centroid for both mice, across the session. The implant key-points are always near mouse0 and there are no identity swaps. **f**, The actual implant-key-point to implant-skeleton-point distance for mouse 0, across the session, is lower than the distance to the centroid of the nose ellipsoid.

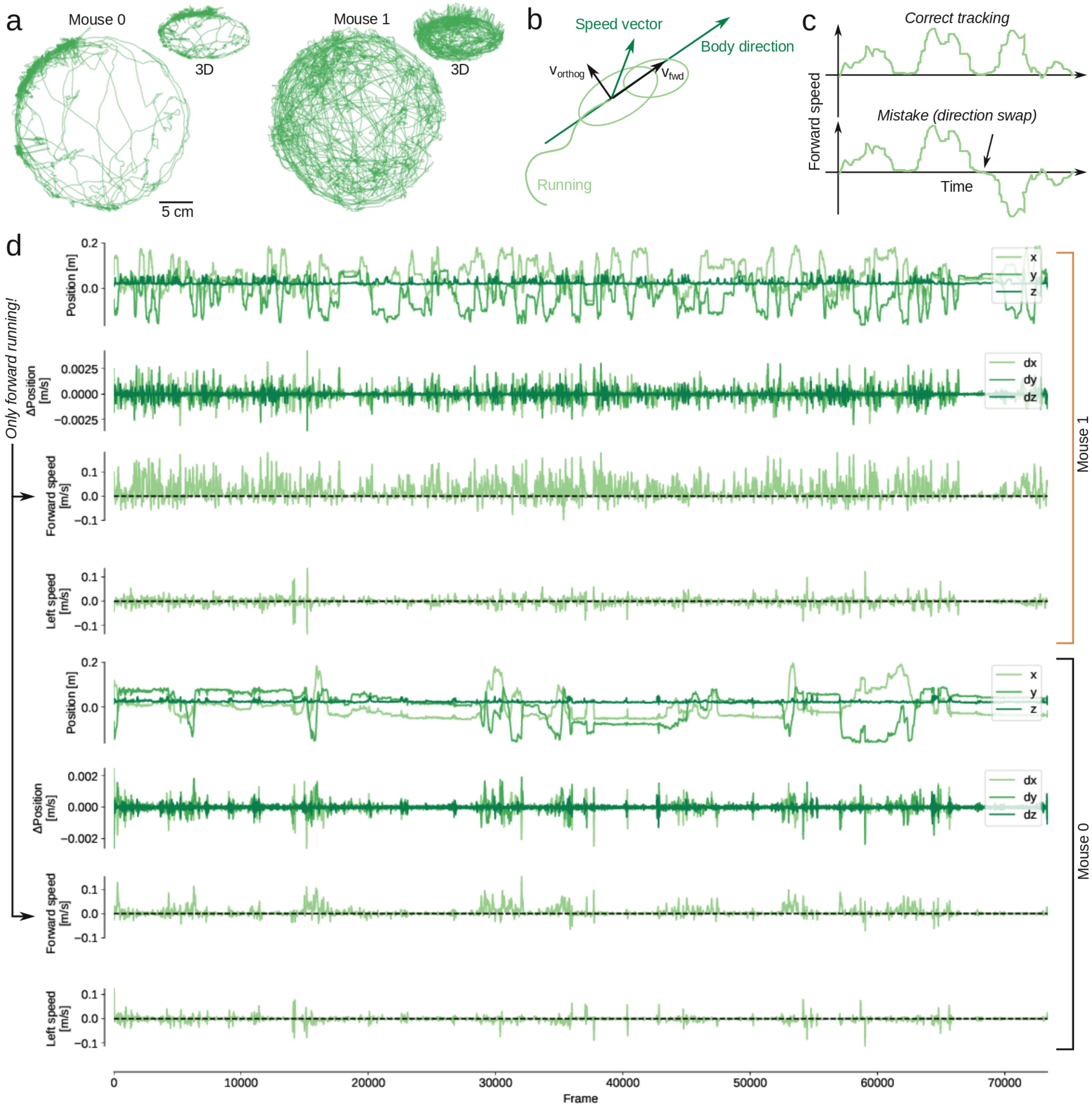


**Supplementary Figure 14. Calculation of movement speed in egocentric coordinates. a**, Running behavior of the two mice (centroid of the hip ellipsoid) across the behavioral session, shown in 2D (top-down view) and 3D. **b**, Running speed decomposed into two components, 'forward speed' (v_fwd, projected onto the orientation of the hip ellipsoid) and 'left speed' (v_orthog, the orthogonal component). **c**, In correct tracking (top row), running bouts will have positive forward speed. If there is a mistake in the tracking (bottom row), such that the mouse body model has switched direction, the mouse will appear to be 'running backwards'. **d**, Top to bottom: The x,y,z-coordinates of the position (c_hip) of the mouse at each tracked frame, the change in position between frames, the forward speed, and the left speed. The four rows are repeated for both mice. There are no direction swap mistakes, and across the whole session, both mice only displayed bouts of forward running (confirmed by visual inspection of raw video).

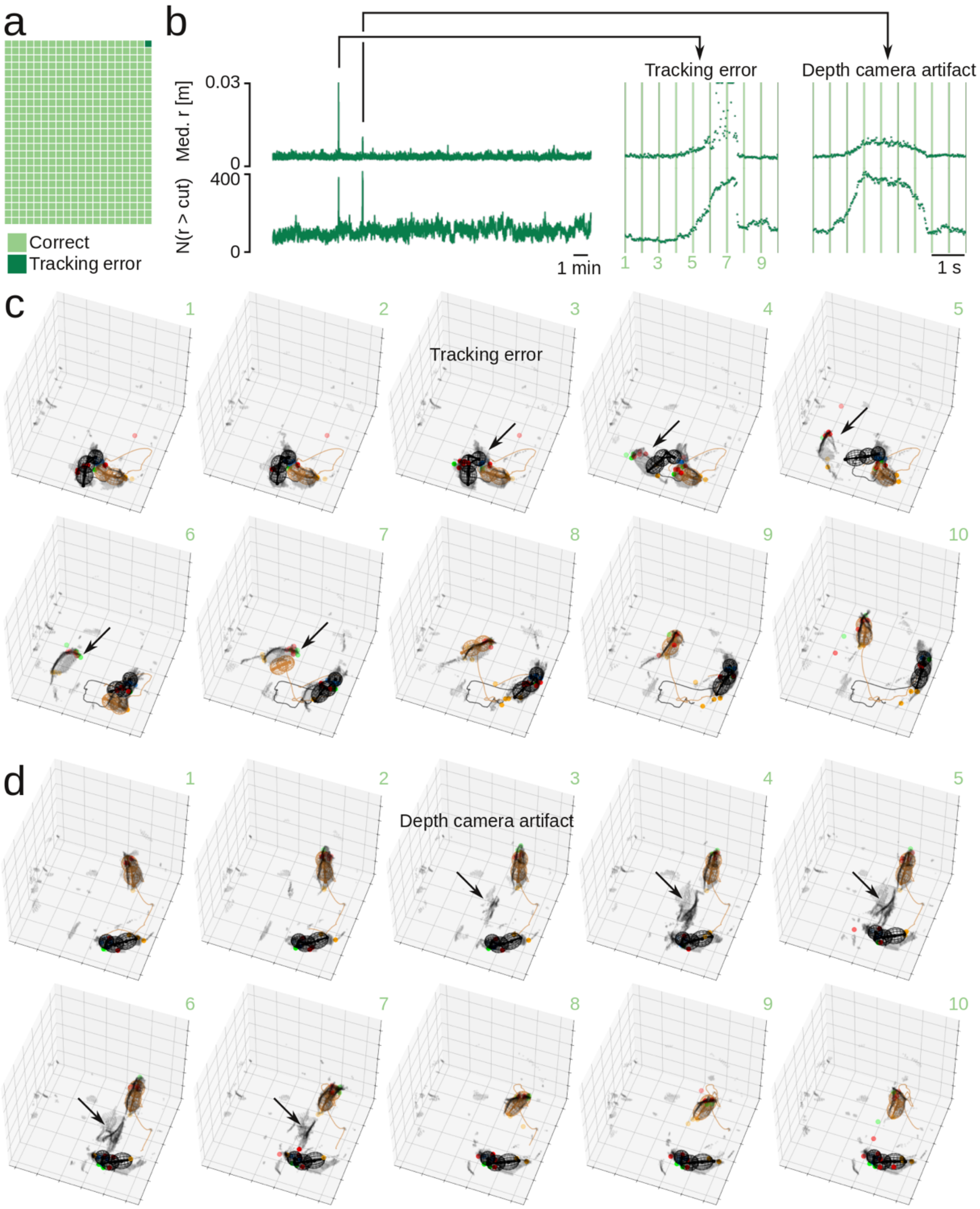


**Supplementary Figure 15. Manual error checking. a**, By manual inspection of 500 frames, we detected one tracking error. **b,** Median point-cloud residual (top) and number of point-cloud points with a residual larger than the cutoff (bottom, cut = 0.03 m) across an example 21 min recording. These traces show two anomalies: One tracking error (around frame 17000, the error we also detected by manual inspection of the 500 frames) and one depth camera artifact (tracking was fine, but a ghostly artifact showed up in the point-cloud for few a seconds. Due to the of the robust loss function, tracking was not distorted by the artifact). **c,** Ten example frames showing the tracking error (0.5 s between frames, indicated by vertical lines in panel b). Note that after the error, the particle filter quickly recovers to correct tracking again. **d,** Ten example frames showing the depth camera artifact.

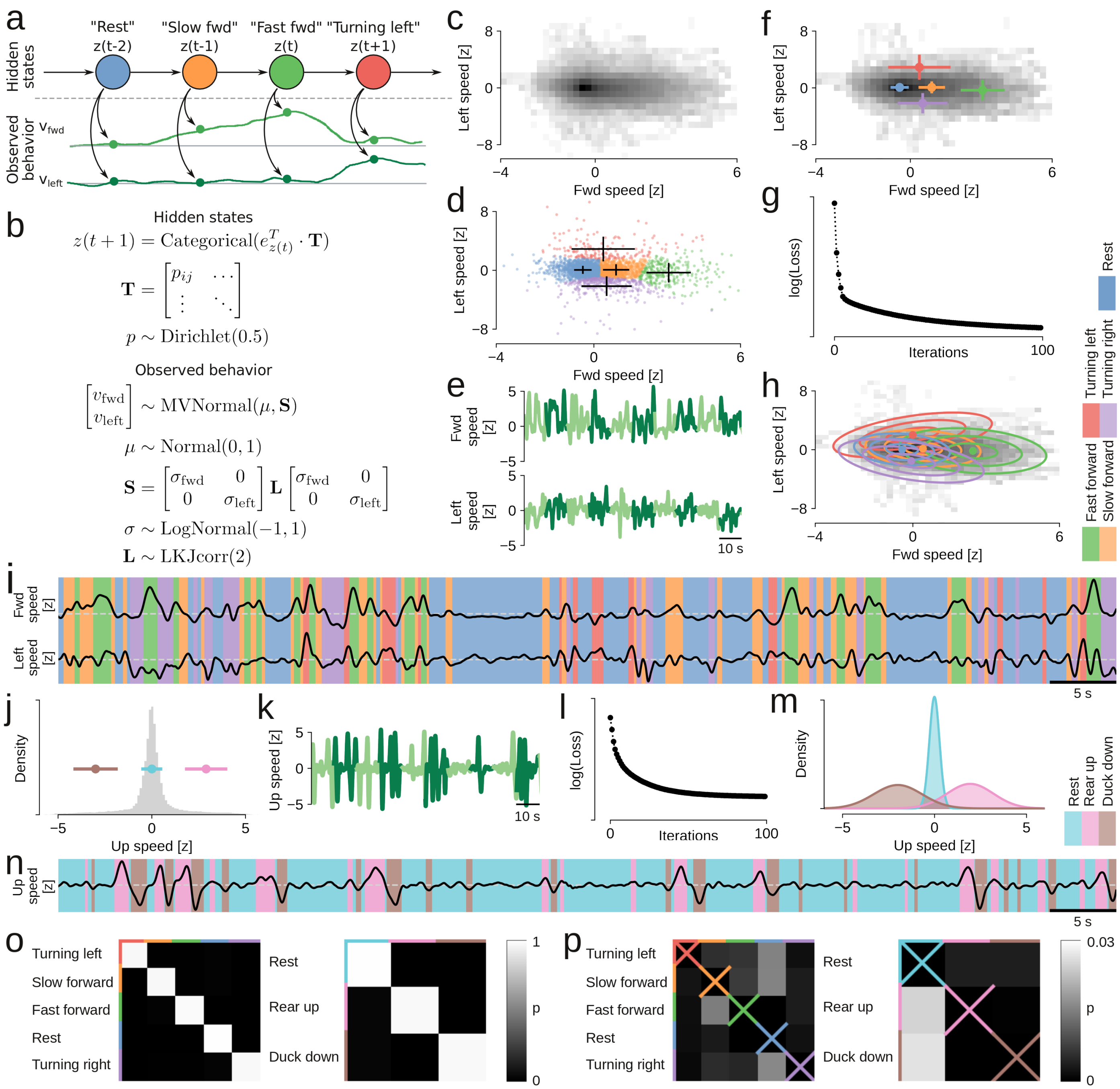


**Supplementary Figure 16. Bayesian modeling and automatic classification of behavioral states. a**, Generative model fit to the running behavior to automatically classify behavioral states. The model is a hidden Markov model with discrete latent states (circles), and each state emits a forward speed and a left speed, drawn from a two-dimensional gaussian distribution with a full covariance matrix. **b**, The generative model expressed as equations, showing Bayesian priors for estimating the parameters. **c**, Joint distribution (on a log-scale) of the forward speed and left speed, for both mice, across an entire behavioral session. **d**, Initial position for the variation inference, for the model of forward and left speed. Crosses, cluster centers and standard deviations (calculated independently for fwd/left speed) for clusters assigned by k-means clustering into 5 clusters. Dots, individual samples of fwd/left speed (colors indicate clusters, every 50th sample is show). **e**, Bayesian model was fitted to a subset of the data (5 mins), split and run in parallel on 10-s sequences. The plot shows example 10-s sequences. **f**, Joint distribution (on a log-scale) of the forward speed and left speed, for the training data, overlaid with the cluster centers and standard deviations from all data (i.e., from **d**). Training data cover same velocity space as the whole session (compare with **a**). **g**, Convergence plot showing the decrease in model loss (increased evidence lower bound) across iterations for the training data. **h**, Locations and covariance ellipsoids (indicating three standard deviations) for the gaussian emission distributions associated with the five latent states, after model fitting. The five clusters are easily interpretable, and the labels are shown on the right. **j**, Initial position for the variation inference for the up speed. Distribution of the up speed (grey bars), as well as the center and standard deviation of three clusters (colored bars and dots), assigned by k-means clustering. **i**, Automatically-assigned states (by maximum a posterior probability) to an example sequence of forward and left speed. **k,l,m,n**, same as **e,g,h,i**, but for the model fitting of the emission gaussians (in one dimension) of the up speed. **o**, Transition probabilities between latent states, for both forward/left speed and up speed models. The sample rate is 60 frames/s, so – since behavioral states are longer than that – the self-transition probabilities (diagonals) are very high. **p**, As **o**, but without showing the self-transition probabilities (the diagonals, crossed out). These matrices have understandable structure. For example, in the left matrix, the most likely transition from “rest” is to “slow forward”. From “slow forward”, the mouse is likely to transition to “turning left”, “fast forward” or “turning right”. It is very unlikely to transition directly from “fast forward” to “rest” or from “turning left” to “turning right”. From the right matrix, we can see that it is unlikely to transition directly from “rear up” to “rear down”, it is more probable to have a period of “rest” in between.

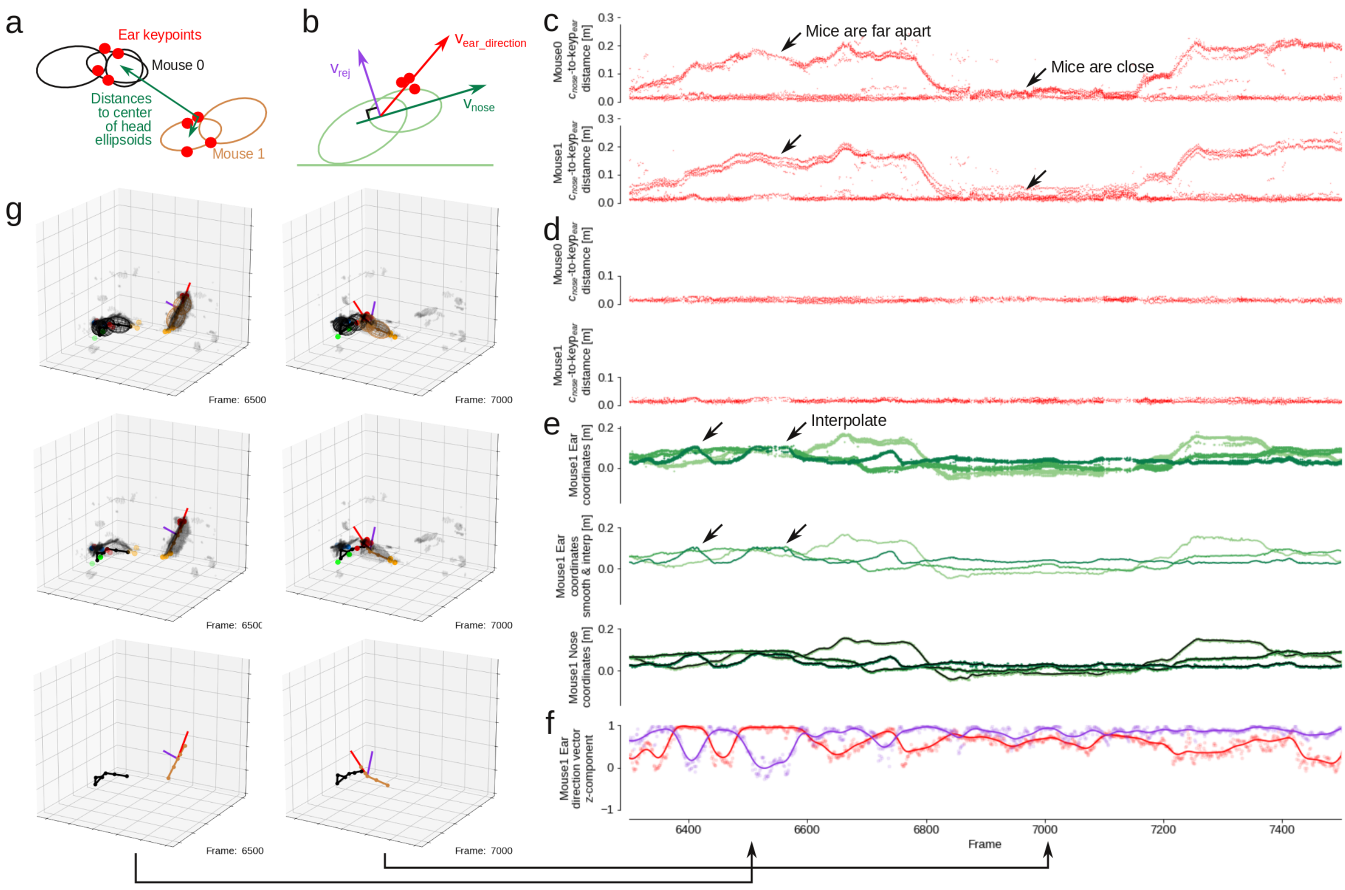


**Supplementary Figure 17. Estimation of 3D heading direction in the partner animal, part I. a**, We use the 3D position of the ear keypoints to determine the 3d head direction of the partner animal. We assign the ear keypoints to a mouse body model by calculating the distance from each keypoint to the center of the nose ellipsoid of both animals. **b**, To estimate the 3D head direction, we calculate the unit rejection (v_rej) between a unit vector along the nose ellipsoid (v_nose) and a unit vector from the neck joint (c_mid) to the average 3D position of the ear keypoints that are associated with that mouse (v_ear_direction). **c**, The distance from all ear keypoints to the center of the nose ellipsoid, for both mice, for an example portion of the recording session. **d**, The distance from ear keypoints to the center of the nose ellipsoid, only showing the keypoints that we estimate to be associated with each mouse. **e**, Estimated mean 3D position of the ear keypoints associated with the partner animal ('Mouse 1'). Top to bottom: Raw 3D position of all keypoints, mean position using linear interpolation, smoothed with a Gaussian kernel ($\sigma = 3$ frames). **f**, The z-component of v_ear_direction and v_rej. The z-component is high, indicating that the ears are on the dorsal side of the head ellipsoid. When the mouse is running on the ground, both v_ear_direction and v_rej have high z-components (marked with rightmost arrow), but when the mouse is rearing and tilting the head backwards, v_rej will be more in the xy-plane, and have a low z-component (marked with leftmost arrow). **g**, The 3D body positions, of the frames indicated by arrows in panel **f**.

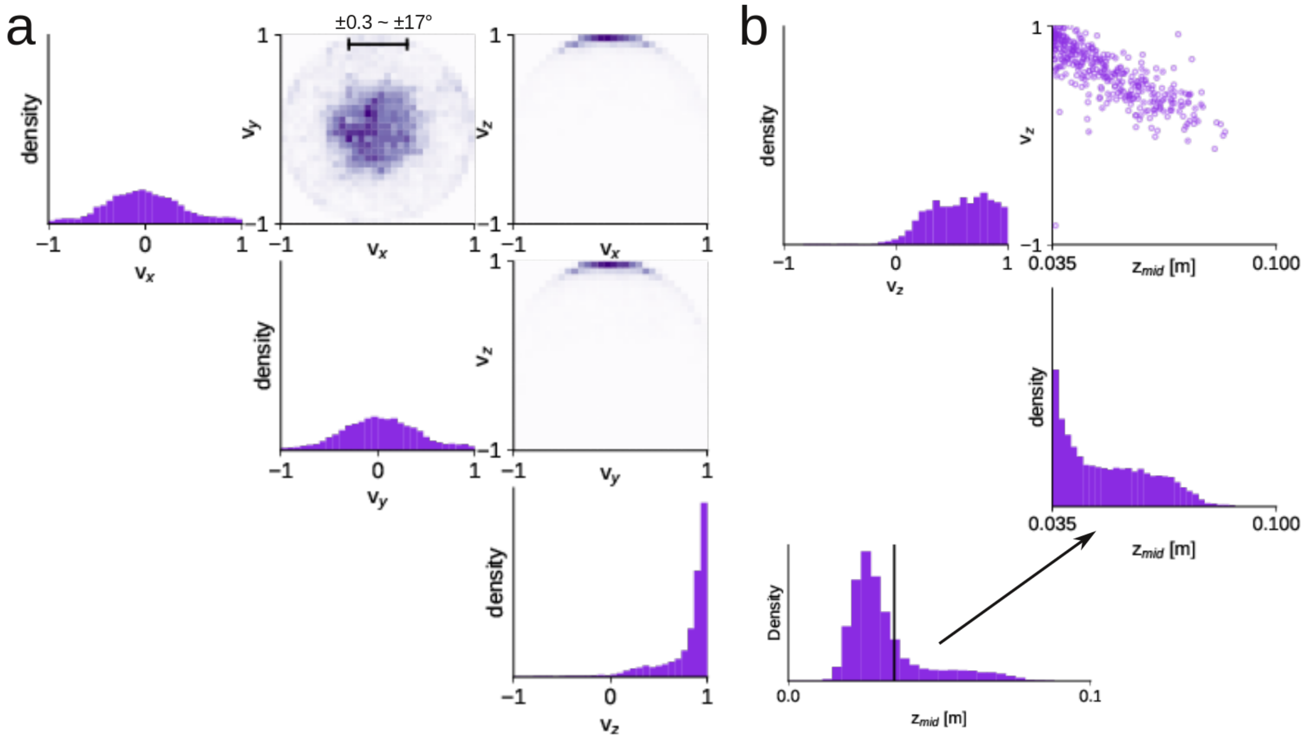


**Supplementary Figure 18. Estimation of 3D heading direction in the partner animal, part II. a**, The joint distributions of the components of v_rej shows that mouse mostly keeps the ears horizontal, rarely tilting the head more than 17 degrees towards the left or right. The z-component is mostly close to 1 (pointing straight up), but sometimes smaller, closer to 0 (meaning that the nose is pointing up towards the sky). **b**, We can examine the details of the 3D head direction behavior. For example, we can monitor the head direction, when the z-coordinate of the neck (z_mid) is high (i.e., when the mouse is rearing). Here we find a clear negative correlation between the z-component of v_rej and z_mid, which matches the visual inspection of the videos: When the mouse rears up or climbs up against the walls of the transparent social arena, the head tilts back to extend the nose upwards.

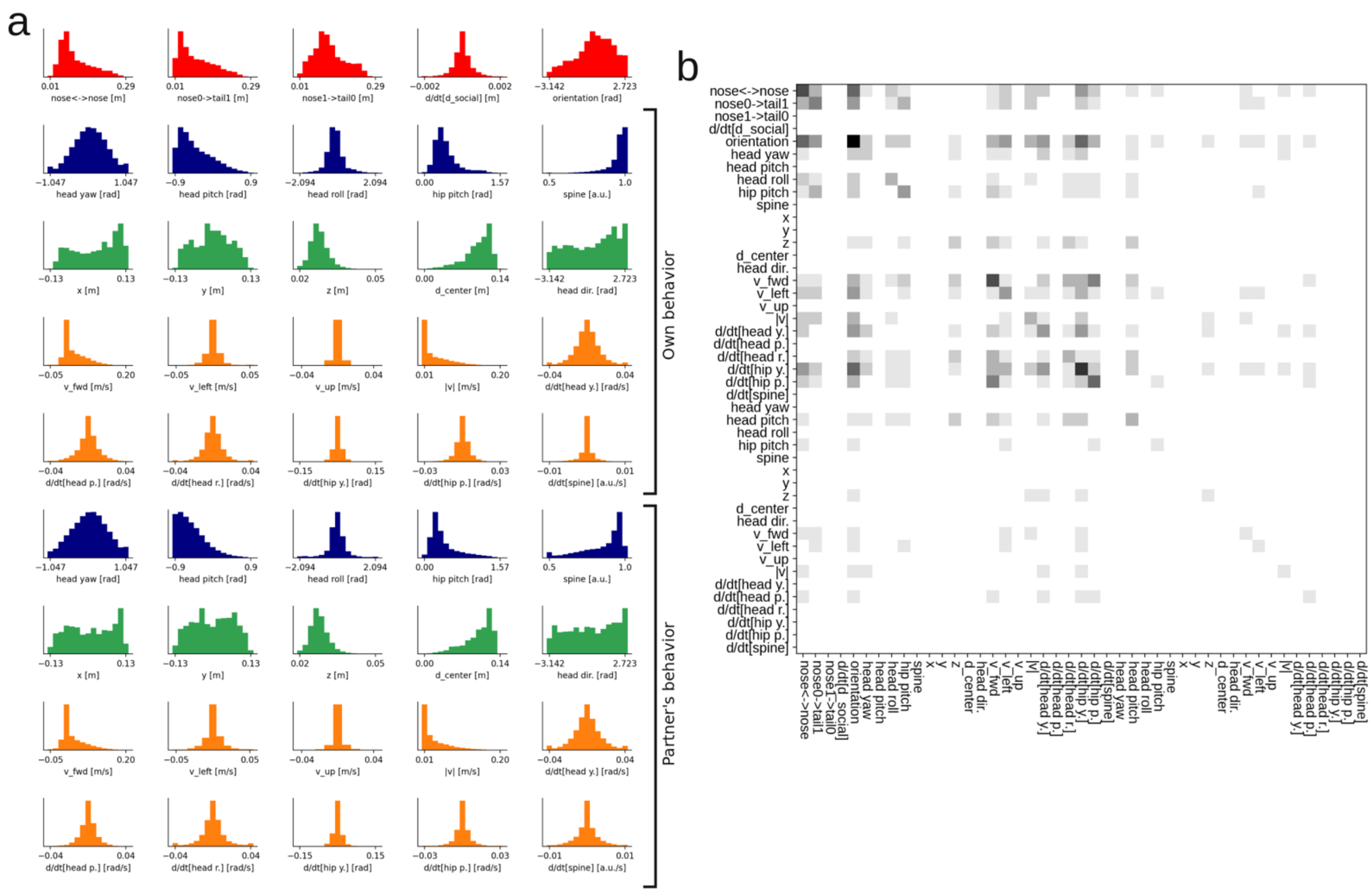


**Supplementary Figure 19. Details of 3D social behavior. a**, Distribution of the behavior occupancy in the bins of the GLM model, for both the social features, the 'own body' features and the 'partner body' features. **b,** The 'co-encoding matrix' of the neural population: The grayscale color in i'th and j'th bin in the heatmap indicates the number neurons that encode both feature i and j, shown here with the full variable names on the matrix axes.

#### Supplementary Videos and Legends

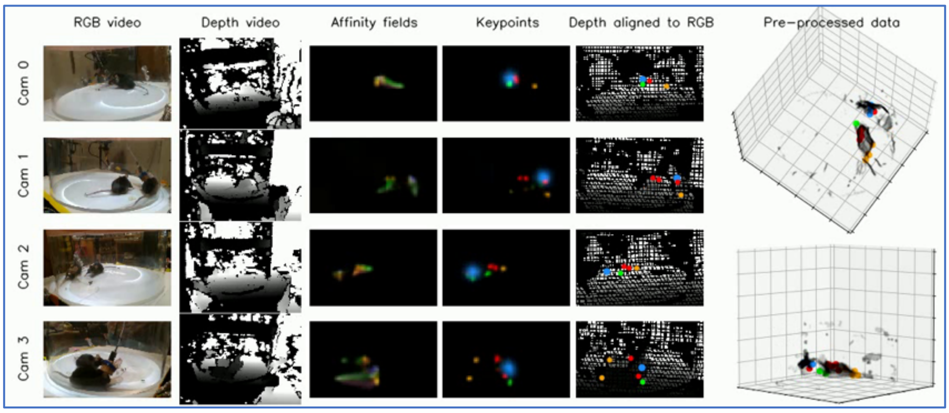


**Supplementary Video 1. Pre-processing pipeline.**

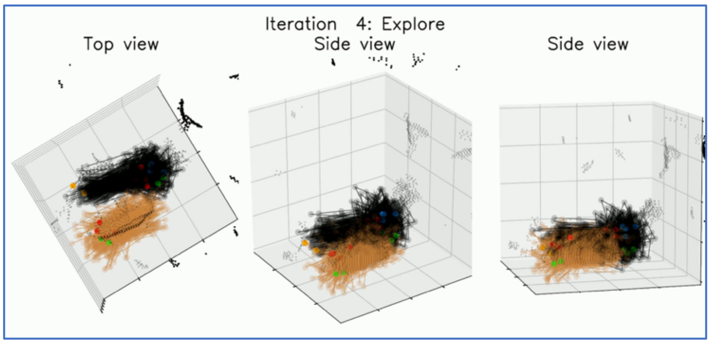


**Supplementary Video 2. Particle filter behavior.**

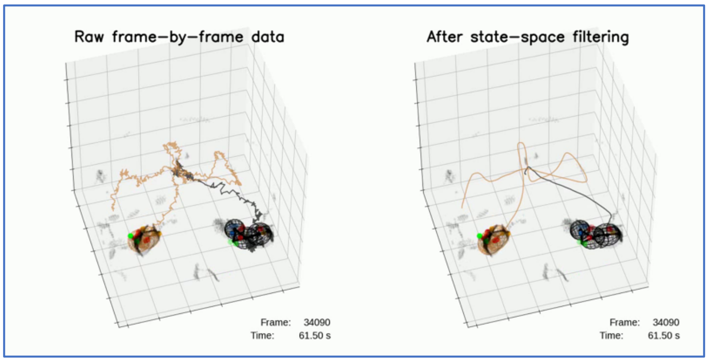


**Supplementary Video 3. State-space filtering.**

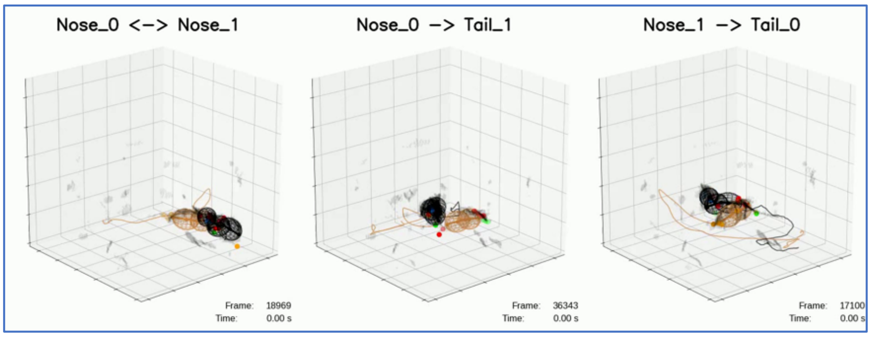


**Supplementary Video 4. Social events.**

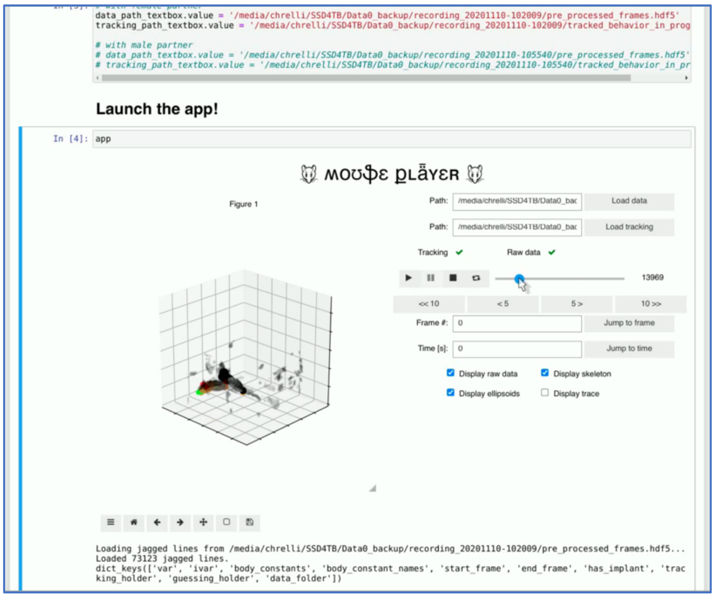


**Supplementary Video 5. MousePlayer.**
